# HTLV-1 antisense transcription is promoted by increased SP1 binding at 3’-LTR G-Quadruplexes

**DOI:** 10.1101/2025.10.11.681801

**Authors:** Emanuela Ruggiero, Irene Zanin, Beatrice Tosoni, Sara N. Richter

## Abstract

The human T-cell lymphotropic virus type 1 (HTLV-1) is a highly oncogenic delta-retrovirus. It presents 5’- and 3’-long terminal repeats (LTR) that are enriched in putative G-quadruplex (G4)-forming sequences. G4s are non-canonical nucleic acid structures that regulate key biological processes in both human and viral genomes.

We here investigated the presence and functional role of G4s within the HTLV-1 3’-LTR, which governs the antisense transcription of the viral bZIP factor (HBZ), the main responsible for T-cell transformation. We identified seven highly conserved sequences that folded into two-layer G4s in both single- and double-stranded DNA in vitro. We demonstrated G4 folding in infected cells by chromatin immunoprecipitation and showed SP1 enrichment at the 3’-LTR G4s. We showed that G4 stabilization with a ligand enhances antisense transcription by promoting recruitment of SP1.

Our findings unveil a G4-mediated regulatory mechanism sustaining HTLV-1 antisense transcription and provide new insights into the complex interplay between the HTLV-1 genome and host cellular factors, contributing to our understanding of retroviral replication strategies to be exploited as new therapeutic targets.

## INTRODUCTION

The human T-cell lymphotropic virus type 1 (HTLV-1) is a highly oncogenic delta-retrovirus (delta-RV) associated with two major pathologies: adult T-cell leukemia (ATL), an aggressive T-cell malignancy (Nosaka & Matsuoka, 2021), and HTLV-1-associated myelopathy/tropical spastic paraparesis (HAM/TSP), a progressive neuroinflammatory condition (Ahmadi Ghezeldasht *et al*, 2024). Among oncoviruses, HTLV-1 exhibits exceptional carcinogenic potential, with 5-10% of infected individuals developing cancer. Current epidemiological studies estimate a global infection burden of approximately 15 million people (Branda *et al*, 2025; Wang *et al*, 2024). Given the limited efficacy of available treatments and the consequent poor prognosis, a deeper understanding of viral pathogenesis is urgently needed.

The HTLV-1 genome consists of two copies of positive-sense, single-stranded RNA that, upon host cell entry, is reverse-transcribed into double-stranded DNA and integrated into the host genome, where it either remains transcriptionally silent or becomes activated, but cannot be eradicated. The provirus is flanked by two identical long terminal repeats (LTRs), critical for viral replication, composed of unique 3′ (U3), repeated (R), and unique 5′ (U5) regions (Ma *et al*, 2016). The 5’-LTR acts as a promoter for structural, accessory, and regulatory viral genes, whereas the 3’-LTR drives transcription of the HTLV-1 bZIP factor (HBZ) from the antisense strand (Matsuoka & Mesnard, 2020). HBZ plays a pivotal role in maintaining viral latency and promoting T-cell transformation, potentially leading to ATL development. Notably, ATL cells often harbour extensive mutations or deletions in the 5’-LTR, while the *hbz* gene remains conserved and expressed, underscoring its critical role in leukemogenesis (Matsuoka & Mesnard, 2020). Despite its significance, the mechanisms governing *hbz* transcription remain poorly elucidated.

Previous investigations have revealed that most RV LTRs are characterized by high guanine (G) content, enabling the formation of alternative secondary structures known as G-quadruplexes (G4s) (Ruggiero *et al*, 2019; Perrone *et al*, 2017; Kledus *et al*, 2025). G4s form when at least four G-stretches are present, allowing four Gs to establish Hoogsteen-type H-bonds and the resulting tetrads self-stack, forming the G4 structure (Spiegel *et al*, 2020). G4 stability is influenced by several factors, including the number of tetrads, the length and composition of loops connecting the G-tracts, interacting proteins and the cellular environment (Kosiol *et al*, 2021; Spiegel *et al*, 2020). G4s are known to regulate key biological processes both in humans (Spiegel *et al*, 2020; Kosiol *et al*, 2021; Robinson *et al*, 2021) and in viruses (Ruggiero & Richter, 2020; Abiri *et al*, 2021), where their targeting has provided promising antiviral effects (Ruggiero & Richter, 2022). In RVs, putative G4-forming sequences (PQSs) are particularly abundant at the LTR level, with delta-RV showing the highest density (Ruggiero *et al*, 2019). The exploration of G4 structures and their functional significance in retroviral biology has primarily focused on the human immunodeficiency virus type 1 (HIV-1), the etiological agent of the acquired immunodeficiency syndrome (AIDS). In HIV-1, G4s located within the 5’-LTR have been shown to modulate transcription through interaction with cellular proteins, and their stabilization by specific ligands hinders virus propagation (Perrone *et al*, 2015, 2014; Tosoni *et al*, 2015; Ruggiero *et al*, 2022a). In contrast, similar investigations in HTLV-1 remain limited.

In this study, we investigated the presence and functional role of G4s in the HTLV-1 LTRs, focussing on the 3’-LTR G4s and their potential involvement in regulating antisense transcription. We identified and characterized seven highly conserved PQSs in vitro and demonstrated their folding within chromatin in HTLV-1-infected cells. Using the well-established G4 ligand Pyridostatin (PDS), we showed that G4 stabilization increases *hbz* expression, revealing a previously unrecognized mechanism of transcriptional regulation. These findings provide novel insights into HTLV-1 gene expression and unveil new mechanisms in retroviral pathogenesis.

## RESULTS

### The HTLV-1 LTRs are enriched in highly conserved 2-tetrad G4s

Building on our previous observations of high PQS density in delta-RVs, particularly within the LTR regions (Ruggiero *et al*, 2019), we conducted a comprehensive bioinformatic analysis to identify G4-forming motifs in the HTLV-1 provirus LTR. Using the Quadbase2 algorithm (Dhapola & Chowdhury, 2016), we screened both sense and antisense strands, applying three levels of stringency: (i) High, requiring four GGG-tracts and loop length up to 12 nucleotides, (ii) Medium, allowing one 1-nucleotide bulge within one G-tract (Mukundan & Phan, 2013) and loop length up to 12 nucleotides, and (iii) Low, based on four GG-tracts connected by short loops (1-7 nucleotides). The High and Medium criteria identify G4s with at least 3 tetrads, while the Low group includes potentially less stable 2-layer G4s. Our analysis identified seven sequences, designated HTLV-1a-g, located in the reverse strand, all meeting the Low-stringency criteria, thus suggesting a potential prevalence of 2-tetrad G4s within the HTLV-1 promoter (Fig 1A). These GG-based PQSs span the entire LTR, with HTLV-1a-c located in the U3, HTLV-1d-f in the R and HTLV-1g in the U5. Notably, HTLV-1 sense transcription initiates at the U3, just upstream of the TATA box (Fauquenoy *et al*, 2017). On the other hand, the antisense promoter is TATA-less, therefore it includes several initiation sites distributed along the entire 3’-LTR (Manghera *et al*, 2017). The broad distribution of PQSs along the 3’-LTR may therefore be functionally relevant to the regulation of the antisense transcription.

**Figure 1.**
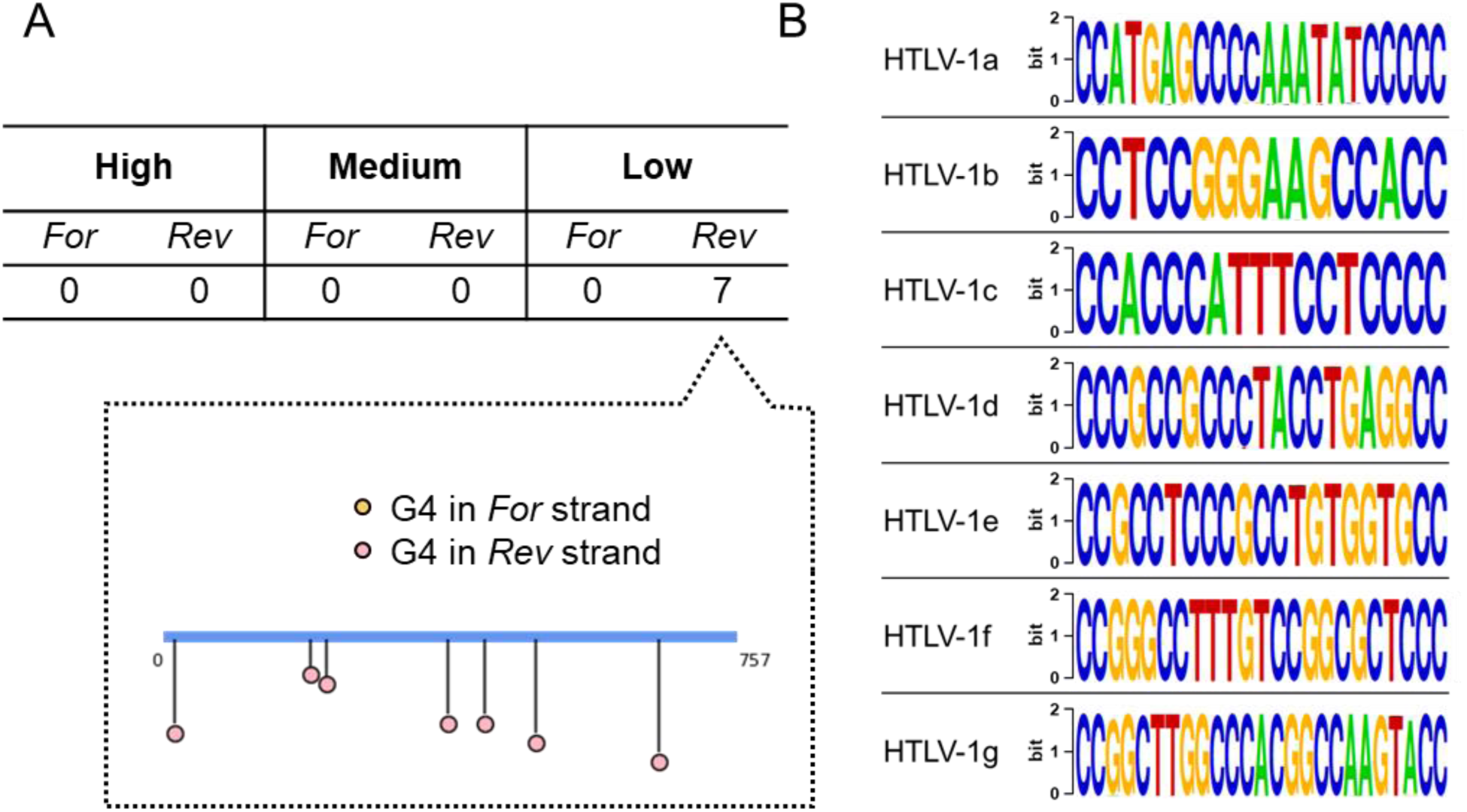
Bioinformatic G4 analysis of HTLV-1 LTR. A) Number of PQSs identified with the Quadbase2 software, classified by G4 search parameters applied to both the forward (*For*) and reverse (*Rev*) strands. Applied prediction criteria: High, (G_3_L_1-12_)_3_G_3_; Medium (G_3_L_1-12_)_3_G_3_ comprising 1-nucleotide bulge within one G-tract; Low, (G_2_L_1-7_)_3_G_2_. G indicates the guanines in each G-tract while L indicates the nucleotides forming the loops. The lollipop graph indicates the distribution of the identified PQSs throughout the LTR region. B) Base conservation of putative 2-layer G4s. Consensus sequences, reported as Weblogo outputs, were derived from the alignment of 256 HTLV-1 strains.

To further explore the biological relevance of these sequences, we investigated the conservation of the identified PQSs across more than 250 HTLV-1 strains. Despite the high genetic variability typical of RVs, primarily due to error-prone reverse transcription and recombination events during integration (Rethwilm & Bodem, 2013), we observed striking conservation of G residues (Fig 1B). This conservation was especially pronounced in G-tracts involved in putative G4s, consistent with observations in arboviruses (Nicoletto *et al*, 2023), retroviruses (Ruggiero *et al*, 2019; Perrone *et al*, 2017), and herpesviruses (Frasson *et al*, 2019; Biswas *et al*, 2018). Furthermore, high conservation was also observed in the loop nucleotides, which are generally implicated in protein recognition (Lago *et al*, 2017; Nagatoishi & Sugimoto, 2012). These findings reinforce the hypothesis that G4 structures play critical roles in viral life cycles and suggest positive selection of these elements during viral genome evolution.

We next evaluated the folding propensity of the bioinformatically predicted 2-layer G4s using circular dichroism (CD) spectroscopy. All but one tested oligonucleotides demonstrated the ability to adopt G4 structures, albeit with diverse topologies (Fig 2A). Specifically, HTLV-1a and HTLV-1c exhibited a major positive peak at λ = 260 nm and a minimum at λ = 240 nm, indicative of a predominant parallel G4 conformation (Kypr *et al*, 2009). The HTLV-1d sequence formed a mostly antiparallel G4, as evidenced by a positive peak at a higher wavelength (290 nm) and a minimum around 260 nm (Kypr *et al*, 2009). HTLV-1b adopted a hybrid conformation (del Villar-Guerra *et al*, 2018), whereas HTLV-1e and −1f showed broad, less-defined peaks in the 270–280 nm range, likely indicating the presence of multiple conformations in solution. HTLV-1g, the only outlier, displayed a spectrum consistent with predominantly unstructured DNA (Fig 2A). We further assessed HTLV-1 LTR G4s stability by monitoring structure unfolding at increasing temperatures, from 20 to 90°C (Appendix Fig S1-S2). Melting temperatures (T_m_), defined as the temperature at which half of the G4 structure is unfolded, ranged from 43°C to > 90°C (Table 1), suggesting that the oligonucleotides likely maintain their folded conformation under physiological conditions. For HTLV-1g, T_m_ could not be determined, likely due to the presence of multiple unfolding transitions, supporting the absence of a stable G4 structure (Appendix Fig S2G).

**Figure 2.**
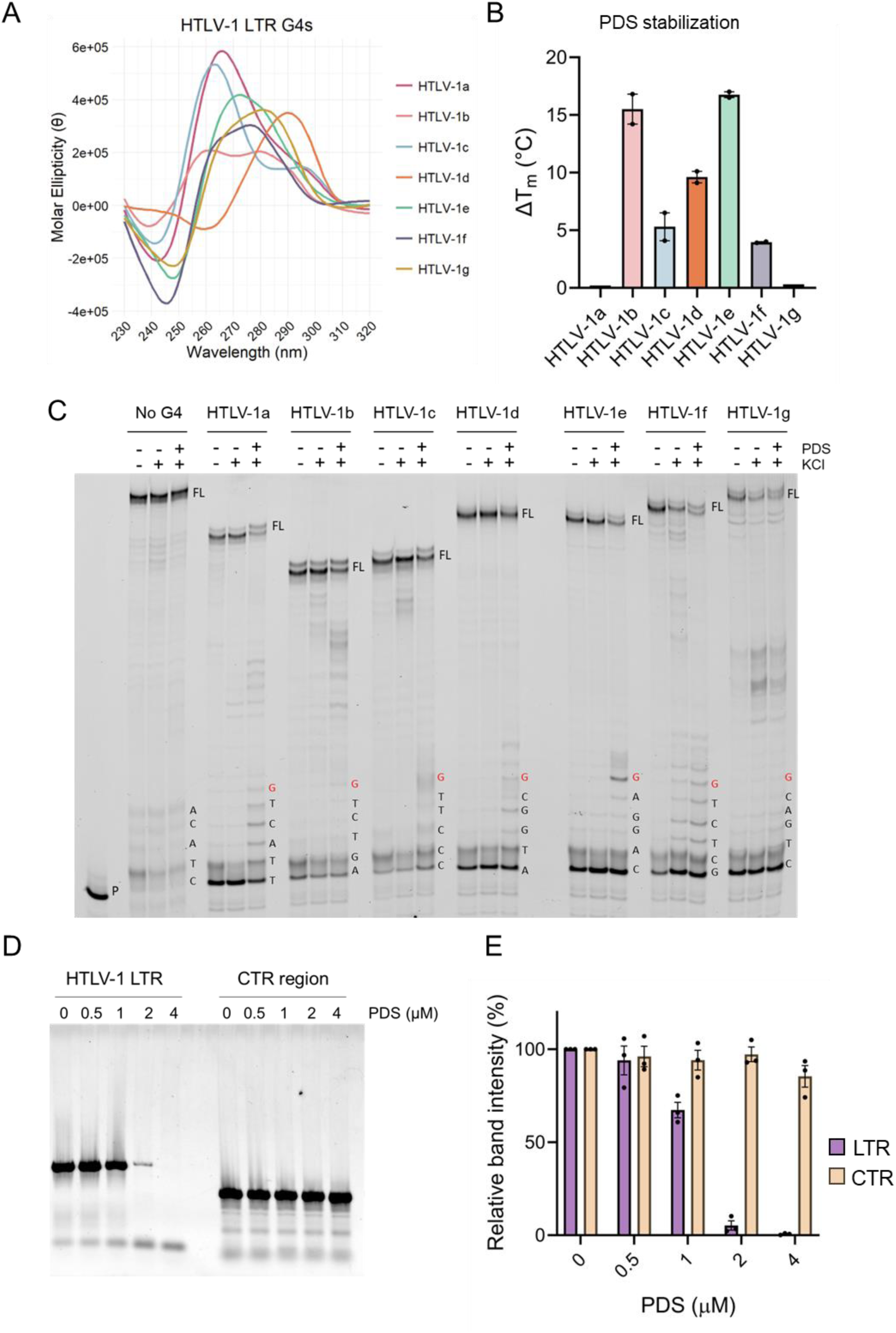
*In vitro* characterization of HTLV-1 LTR G4s. A) CD spectra of HTLV-1 LTR G4s performed in lithium cacodylate/KCl buffer at 3 µM final concentration. Molar ellipticity was measured at 20°C and reported as deg x cm^2^ x dmol^-1^. B) Differences in melting temperature (ΔT_m_) induced by PDS treatment (4-molar excess) and measured by thermal unfolding analysis. For each oligonucleotide, the ΔT_m_ value is reported as mean ± s.e.m. of n = 2 experiments. C) Representative gel of the *Taq* polymerase stop assay performed on HTLV-1 LTR G4s (n=2). Templates were amplified by *Taq* polymerase in the absence or presence of KCl, and KCl combined with 1µM PDS. A non-G4 forming template (No G4) was used as negative control. P indicates the unreacted labeled primer; FL indicates the full-length product. The first G of the sequence involved in G4 formation is highlighted in red. D) PCR stop assay on gDNA extracted from MT-2 cells performed in the absence and presence of increasing concentrations of PDS. The CTR region indicates a non-G4 forming genomic region used as negative control. E) Quantification of the gel bands shown in panel (D). Intensity of the bands from the PCR reactions was normalized on the untreated sample. Violet bars indicate amplification of the HTLV-1 3’-LTR, yellow bars indicate the control region. Data are reported as mean ± s.e.m. (n = 3).

**Table 1.** Melting temperatures obtained by CD thermal unfolding in the absence and presence of PDS.

|  | T<sub>m</sub> Oligo (°C)<sup>a</sup> | T<sub>m</sub> Oligo + PDS (°C)<sup>a</sup> |
| --- | --- | --- |
| HTLV-1a | >90 | >90 |
| HTLV-1b | 54.4 ± 0.1 | 69.9 ± 1.1 |
| HTLV-1c | 58.9 ± 0.3 | 64.2 ± 1.5 |
| HTLV-1d | 43 ± 0.2 | 53.6 ± 0.3 |
| HTLV-1e | 58.3 ± 0.1 | 75.0 ± 0.3 |
| HTLV-1f | 59.8 ± 0.6 | 63.8 ± 0.6 |
| HTLV-1g | ND | ND |

<sup>a</sup>Values are reported as mean ± s.e.m. of two independent experiments.

### PDS stabilizes HTLV-1 LTR G4s

We subsequently investigated the impact of PDS on HTLV-1 LTR G4 stability. PDS was selected due to its well-established role as a potent G4 stabilizer with promising antiviral properties (Ruggiero & Richter, 2022). CD measurements revealed a general increase in thermal stability upon PDS binding, with T_m_ shifts (ΔT_m_) ranging from approximately 4°C for HTLV-1f to 17°C for HTLV-1e (Fig 2B, S2 and Table 1). Notably, both HTLV-1e and −1f sequences showed PDS-mediated stabilization, supporting the coexistence of G4 and non-G4 conformations in solution, as suggested by their CD spectra recorded in the absence of the ligand (Fig 2A) (Ruggiero *et al*, 2019). HTLV-1a G4 already exhibited high intrinsic stability under the assay conditions, precluding reliable detection of further stabilization by PDS (Appendix Fig S2A). In contrast, HTLV-1g showed no conformational changes upon PDS treatment, confirming that it does not adopt a G4 arrangement in the tested conditions.

The effect of PDS on HTLV-1 LTR G4s was further evaluated using a *Taq* polymerase stop assay (Fig 2C). This technique exploits the ability of folded G4s to stall polymerase progression, thereby resulting in truncated DNA products (Ruggiero *et al*, 2022b). HTLV-1 LTR G4 sequences were employed as templates, and polymerase progression was monitored under three conditions: absence and presence of potassium to promote G4 formation, and presence of both potassium and PDS. A random, non-G4-forming sequence was included as negative control. Minimal differences were observed between potassium-free and potassium-containing conditions, likely due to the lower stability of 2-layer G4s. However, all tested sequences but HTLV-1g exhibited substantial polymerase stalling upon PDS treatment, reflected by a marked decrease in full-length product synthesis and the appearance of truncated products. These stalling events occurred at or near the first guanine involved in G4 folding (highlighted in red in Fig 2C) (Ruggiero *et al*, 2022b), confirming G4 formation. For the HTLV-1g sequence, stop bands were also detected, though at positions incompatible with G4 formation, in agreement with the CD data. The negative control template showed no polymerase stalling, supporting the specificity of the effect.

To investigate G4 formation within the context of the integrated provirus, we performed a PCR stop assay on genomic DNA (gDNA) extracted from the HTLV-1-infected MT-2 cell line. PCR primers flanking the full-length 3’-LTR were used to amplify this region, which includes all PQSs identified in our study, while a non-G4-forming genomic region served as a control (Fig 2D). Treatment with increasing concentrations of PDS resulted in a strong, dose-dependent reduction in full-length amplicon intensity (Fig 2E), indicative of polymerase stalling. Amplification of the control region remained unaffected by PDS, validating the G4-specificity of the observed effect. These results confirm that G4s sufficiently stable to impede polymerase progression in vitro form in the HTLV-1 proviral LTR.

### LTR G4s promote antisense transcription via SP1 recruitment

To further elucidate the role of HTLV-1 G4s located at the 3’-LTR in a chromatin context, we performed chromatin immunoprecipitation (ChIP) on HTLV-1-infected MT-2 cells using the G4-specific antibody BG4 (Maurizio *et al*, 2024). Quantitative PCR (qPCR) was used to assess G4 enrichment at two distinct LTR regions: LTR-1 and LTR-2. The LTR-1 region lies within the LTR and is therefore identical at both proviral termini. This region encompasses the HTLV-1b and HTLV-1c G4s. The LTR-2 region covers only the antisense promoter, including the R-U5 regions of the 3’-LTR terminus and extending into the adjacent gene sequence (Yamagishi *et al*, 2021). GAPDH and HTR-6 loci served as positive and negative controls for G4 formation, respectively (Hansel-Hertsch *et al*, 2016). Samples included BG4-immunoprecipitated DNA, a mock immunoprecipitation without antibody, and the input chromatin for normalization. G4 enrichment for each region was then reported as fold change relative to the negative region. Both LTR regions exhibited significant G4 enrichment, comparable to the positive control (Fig 3A), providing the first direct evidence of G4 folding within the integrated HTLV-1 proviral DNA in cells.

**Figure 3.**
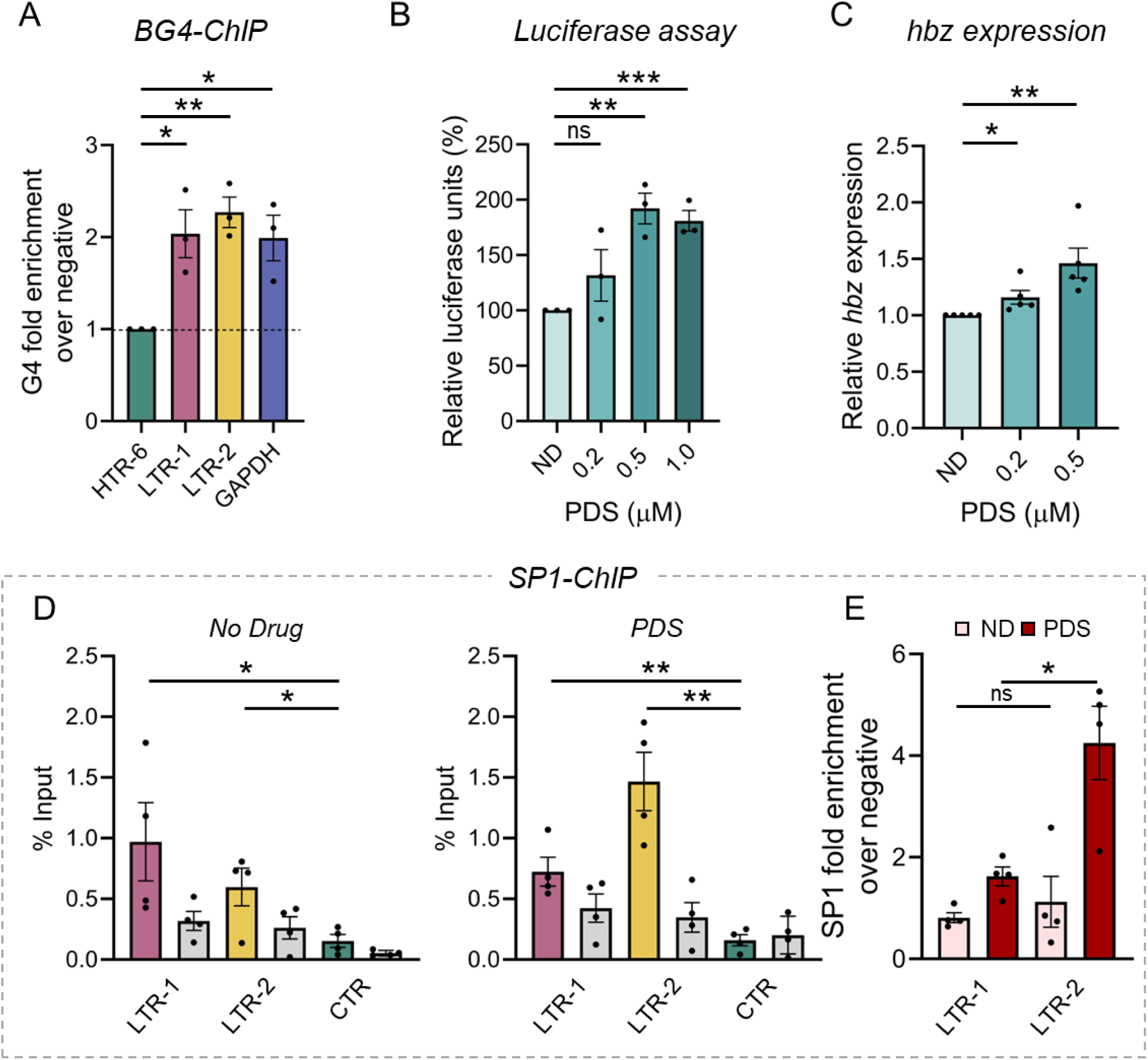
HTLV-1 LTR G4 characterization in cells. A) G4-ChIP-qPCR at the LTR-1 and LTR-2 regions in the chromatin derived from MT2 cells. Results are reported as enrichment over the G4-negative region HTR-6 (dashed line). GAPDH was used as G4-positive control. Data are reported as mean of n = 3 experiments ± s.e.m. Significance levels were calculated using t-test (*p < 0.05, ** p < 0.01). B) Luciferase reporter assay of HTLV-1 LTR antisense promoter activity in the presence of increasing amounts of PDS. Data are reported as mean of n = 3 experiments ± s.e.m. Significance levels were calculated using t-test (ns = p ≥ 0.05, ** p < 0.01, *** p < 0.001). C) *Hbz* transcript levels in MT-2 cells untreated (ND) or treated with PDS and normalized on the *gapdh* housekeeping gene. Data are reported as mean of n = 5 experiments ± s.e.m. Significance levels were calculated using t-test (* p < 0.05, ** p < 0.01). D) SP1-ChIP-qPCR at the LTR-1 and LTR-2 regions in the chromatin derived from MT-2 cells, performed in the absence (left panel) or presence (right panel) of PDS 0.5 µM. Results are reported as the percentage of input for immunoprecipitated (coloured bars) and mock samples (grey bars). CTR indicates a negative control region for SP1 binding. Data are reported as mean of n = 4 experiments ± s.e.m. Significance levels were calculated using t-test (*p < 0.05, ** p < 0.01). E) Fold enrichment of immunoprecipitated samples normalized over the background and the negative region, in the absence (light pink bars) or presence (dark red bars) of PDS 0.5 µM. Data are reported as mean of n = 4 experiments ± s.e.m. Significance levels were calculated using t-test (*p < 0.05, ns = not significant).

Motivated by this finding, we investigated the functional consequences of G4 stabilization on antisense transcription using a bidirectional reporter plasmid. This plasmid harbors two reporter genes under the control of the full-length HTLV-1 LTR arranged in opposite orientations, enabling discrimination between sense (Renilla luciferase) and antisense (Firefly luciferase) promoter activity (Arpin-André *et al*, 2014; Laverdure *et al*, 2016). HEK293T cells were treated with increasing concentrations of PDS to establish a sub-cytotoxic dosage range (Appendix Fig S3A). Following transfection and 24 h PDS exposure, Firefly luciferase activity, representing antisense transcription, increased significantly in a dose-dependent manner (Fig 3B), indicating transcriptional enhancement.

To validate these results in a more physiological setting, HTLV-1-infected MT-2 cells were treated with subtoxic PDS concentrations for 24 h (Appendix Fig S3B) and *hbz* mRNA levels were quantified by qRT-PCR. Consistent with the reporter assay, *hbz* expression increased dose-dependently following PDS treatment (Fig 3C).

Given that antisense promoter activity is largely dependent on SP1 transcription factor binding, which initiates transcription at multiple dispersed sites due to the absence of canonical TATA boxes (Manghera *et al*, 2017; Laverdure *et al*, 2016), we next investigated whether the observed transcriptional enhancement involved increased SP1 occupancy. ChIP-qPCR using an anti-SP1 antibody was performed on MT-2 cells untreated or treated with PDS for 24 h at the highest concentration used in *hbz* expression analysis. We measured SP1 enrichment at LTR-1 and LTR-2, with a known SP1-negative genomic locus as control (Deshane *et al*, 2010). Under basal (untreated) conditions, the two LTR regions showed comparable SP1 enrichment levels, significantly higher than the control (Fig 3D). To note that the signal from LTR-1, being present at both the 5’- and 3’-LTR, represents an average of the signals from the two regions, precluding precise SP1 localization. Upon PDS treatment, SP1 occupancy at the antisense promoter (LTR-2) significantly increased, while enrichment at the LTR-1 region remained unchanged (Fig 3E). This indicates that PDS-enhanced G4 formation facilitates SP1 recruitment at the 3’-LTR, supporting a G4-dependent transcriptional regulation at the HTLV-1 antisense promoter.

## DISCUSSION

Despite its initial discovery in the early 1970s, research on HTLV-1 molecular biology and pathogenesis remains limited. While the pivotal role of *hbz* gene expression and HBZ protein in HTLV-1 pathogenesis is well-established (Ma *et al*, 2016; Baratella *et al*, 2017; Yamada *et al*, 2021), the underlying mechanisms are still incompletely understood. Here, we identified G4s located in the 3’-LTR as novel elements of the antisense transcription machinery, acting by facilitating SP1 transcription factor recruitment.

Our analysis revealed remarkable evolutionary conservation of the identified PQSs within the LTR. Although viruses typically display high genetic variability to evade immune pressure and adapt to environmental conditions, certain genomic regions must remain unaltered, to preserve viral propagation and survival (Holmudden *et al*, 2024). The conservation of LTR G4s across viral isolates thus suggests functional constraint and underscores their potential as therapeutic targets applicable across diverse HTLV-1 strains. These findings also offer new perspectives on the role of structured nucleic acids in retroviral genome evolution.

Interestingly, our data indicate in HTLV-1 3’-LTR G4s are the most relevant as opposed to the 5’-LTR G4s, despite the sequence identity of these regions. This disparity may relate to differential epigenetic regulation: the 5’ sense promoter is frequently hypermethylated in integrated proviruses, leading to transcriptional silencing, whereas the antisense promoter at the 3’-LTR is generally hypomethylated, suggesting the presence of protective mechanisms that preserve *hbz* expression. Indeed, differential methylation at these sites appears necessary for HTLV-1 persistence in vivo (Pluta *et al*, 2020; Kulkarni & Bangham, 2018; Miura *et al*, 2018). Notably, G4s have been demonstrated to protect genomic regions from methylation, especially in open chromatin contexts (Jara-Espejo & Line, 2020; Rauchhaus *et al*, 2022; Niu *et al*, 2025). Our data thus suggest that HTLV-1 LTR G4s may similarly preserve the antisense promoter from methylation, thereby supporting sustained *hbz* transcription.

Structurally, the G4s identified in the HTLV-1 LTR are predominantly characterized by 2-tetrads, generally considered less stable (Kejnovská *et al*, 2021; Heddi *et al*, 2016; Marchand & Gabelica, 2016) and thus often overlooked by computational prediction tools (Hänsel-Hertsch *et al*, 2020; Lam *et al*, 2013). Nevertheless, our chromatin immunoprecipitation assays confirm that these G4s fold in cells (Fig 3A). These findings raise the possibility that more dynamic, transient structures such as the 2-tetrad G4s may be more relevant in viral genome regulation than in eukaryotes. The ability of 2-layer G4s to rapidly fold and unfold (Marchand & Gabelica, 2016) may facilitate interaction with transcription factors, helicases, or ligands, enabling fine-tuned transcriptional control. Indeed, 2-tetrad G4s have been described in the *nef* promoter of the HIV-1 provirus (Perrone *et al*, 2013) and in several RNA viruses including, but not limited to, SARS coronavirus 2 (Gupta *et al*, 2025), influenza virus (Brázda *et al*, 2021) and flaviviruses (Majee *et al*, 2021; Sarkar & Armitage, 2021; Singh *et al*, 2025). This is, to our knowledge, the first direct demonstration of G4 folding within the HTLV-1 integrated (proviral) DNA.

Functionally, we show that the HTLV-1 LTR G4s contribute to the recruitment of SP1 at the 3′-LTR and positively regulate antisense transcription. This is consistent with previous reports demonstrating G4-mediated transcriptional activation in the human genome (Robinson *et al*, 2021). G4 ligands may either enhance or repress gene expression, depending on their impact on protein-G4 interactions. G4 ligands as well as G4-specific antibodies like BG4, may compete with cellular proteins for G4 binding (Spiegel *et al*, 2021; Li *et al*, 2021). PDS has been found to displace the Cockayne Syndrome B protein from G4s located in the ribosomal DNA (Liano *et al*, 2021). Conversely, it binds to EBNA1 mRNA G4 without interfering with nucleolin binding at the same site (Santos *et al*, 2022). Thus, the functional outcome of the G4/PDS/G4-binding protein interplay can be different. In our system, although PDS impaired polymerase progression in vitro, it significantly enhanced transcription in cells, an effect that was G4-dependent and accompanied by increased SP1 recruitment at the 3′-LTR G4 region (Fig 3). These results emphasize the critical influence of the cellular milieu on G4 dynamics and their functional consequences, and further confirm the role of G4s as chromatin marks to recruit transcriptional regulators (Spiegel *et al*, 2021; Lago *et al*, 2021).

Notably, the antisense promoter recruits multiple transcription factors beyond SP1, like JunD, Menin, CEBP and HBZ itself, with JunD acting as a key transcriptional driver (Madugula *et al*, 2022). In this process, HBZ upregulates JunD expression, facilitating formation of a JunD-HBZ complex, which interacts with SP1 bound to the antisense promoter to activate *hbz* transcription (Gazon *et al*, 2012). Our data suggest that G4s mediate the initial recruitment of SP1 to the promoter, thereby facilitating JunD and HBZ binding to promote transcription. Interestingly, a similar pattern has been described at the telomerase gene (*hTERT)* promoter in HTLV-1 infected cells, where HBZ and JunD synergistically enhance *hTERT* transcription through co-recruitment by SP1 (Kuhlmann *et al*, 2007). The *hTERT* core promoter is highly G-rich and capable of folding into G4s encompassing multiple SP1 binding sites (Palumbo *et al*, 2009), suggesting that a similar G4-mediated mechanism may operate at this locus as well.

Given the critical role of *hbz* expression in viral persistence and leukemogenesis, disruption of the G4/SP1 interaction may have therapeutic potential beyond transcriptional regulation, as it could simultaneously impair viral persistence and interfere with oncogenic processes, representing a promising dual strategy against HTLV-1-associated malignancies.

Our study uncovers a previously uncharacterized regulatory network involving LTR-encoded G4s and SP1 in the control of HTLV-1 antisense transcription. The conserved and in vivo folded nature of these two-tetrad G4s highlights their functional significance. These findings advance our understanding on how DNA secondary structures contribute to the regulation of retroviral gene expression and open new avenues for exploring the complex interplay between chromatin architecture and transcriptional control in integrated proviruses.

## MATERIALS AND METHODS

### Oligonucleotides, reagents and cell lines

All the oligonucleotides used in this work were purchased from Sigma-Aldrich and are listed in Appendix Table S1. PDS was obtained from Selleckchem (#S7444). The stock was resuspended in dimethyl sulfoxide at 10 mM final concentration and working solutions were prepared in cell culture medium before use.

MT-2 cells were obtained through the NIH HIV Reagent Program, Division of AIDS, NIAID, NIH: Human T-Lymphotropic Virus Type 1 (HTLV-1)-Infected MT-2 Cells, #ARP-237, contributed by Dr. Douglas Richman. Cells were grown in RPMI medium (Gibco, LifeTechnologies) supplemented with 10% FBS and 1x Penicillin-Streptomycin (Gibco, LifeTechnologies). Human embryonic kidney (HEK293T) cells (ATCC #CRL-3216) were cultured in DMEM medium (Gibco, LifeTechnologies) supplemented with 10% FBS and 1x Penicillin-Streptomycin.

### G4 prediction and base-conservation analyses

Prediction of G4-forming motifs within the HTLV-1 LTR proviral genome was conducted using the QuadBase2 web server (Dhapola & Chowdhury, 2016). Our search parameters encompassed G-tracts composed of three guanines (either continuous or including a single nucleotide bulge) with loops ranging from 1 to 12 nucleotides, as well as G-tracts formed by two contiguous guanines with loops of 1 to 7 nucleotides in length.

To assess the conservation of PQSs in the HTLV-1 LTR, we performed a comprehensive alignment analysis using 256 sequences obtained from PubMed (as of May 2025). The degree of base conservation was visually represented using sequence logos generated by the WebLogo software (Crooks *et al*, 2004).

### Circular Dichroism

Circular dichroism (CD) analysis was performed using oligonucleotides diluted to 3 μM in 20 mM phosphate buffer containing 80 mM KCl. Samples were thermally denatured at 95°C for 5 min, followed by slow cooling to room temperature overnight. CD spectra were recorded using a Chirascan-Plus spectropolarimeter equipped with a Peltier temperature controller, employing quartz cells with 5 mm and 1 mm optical path lengths. Measurements were conducted across a temperature range of 20-90°C, with data acquisition spanning wavelengths from 230 to 320 nm. For experiments involving PDS, oligonucleotides were folded as described above, then the compound was added to a 4-fold molar excess 4 h post denaturation.

All CD data underwent baseline correction and observed ellipticities were converted to mean residue ellipticity (θ) expressed in degree × cm² × dmol⁻¹ (molar ellipticity). CD analyses were performed in duplicate, with values plotted using RStudio software (RStudio 2023.12.0) for windows. Posit team (2025). RStudio: Integrated Development Environment for R. Posit Software, PBC, Boston, MA. URL http://www.posit.co/.

### *Taq* polymerase stop assay

FITC-tagged primer (72 nM) was annealed to template sequences (36 nM) in lithium cacodylate buffer (10 mM, pH 7.4), in the absence and presence of 100 mM KCl. Annealing was achieved by heating the samples to 95°C for 5 min, followed by gradual cooling to room temperature. Where indicated, samples were then incubated overnight at room temperature with PDS (1 μM). Primer extension was performed using AmpliTaq™ 360 DNA Polymerase (2U per reaction; ThermoFisher Scientific, # 4398818) at 37°C for 30 min. Reactions were stopped by ethanol precipitation. The resulting primer-extension products were resolved on a 16% denaturing gel and visualized using phosphorimaging (Typhoon FLA 9000, GE Healthcare, Milan, Italy).

### PCR stop assay

Genomic DNA (gDNA) was extracted from 2×10^6^ cells using the GeneJET Genomic DNA Purification Kit (ThermoFisher Scientific, #K0721) following the manufacturer’s protocol. For the standard PCR amplification of the target regions, 250 ng of gDNA were used as template with DreamTaq DNA polymerase (ThermoFisher Scientific, #EP0701) and incubated with PDS for 4 h at RT before PCR amplification. PCR products were resolved on a 1% agarose gel containing GelRed (Millipore, #SCT122) and electrophoresed at 70 V for 1 hour. Primers used for PCR amplification are listed in Appendix Table S1. PCR band intensity was quantified by ImageQuant TL software and normalized on the untreated sample. Data are reported as mean ± s.e.m. (n = 3).

### Cell viability assay

PDS effect on cell viability was assessed by MTT Cell Growth Kit (Merck, #CT02), according to manufacturer’s instructions. HEK293T and MT-2 cells were seeded in 96-well plates and incubated overnight at 37°C in a humidified atmosphere at 5% CO_2_. Cells were then treated with serial dilutions of PDS, as indicated. 24 h post treatment, 5 mg/ml 3-(4,5-dimethylthiazol-2-yl)-2,5-diphenyltetrazolium bromide solution was added to each well. After 4 h incubation at 37°C and the solubilization of formazan with isopropanol/0.04 N HCl, the absorbance was measured at 620-nm wavelength using the Varioskan LUX Multimode Microplate Reader (ThermoFisher Scientific) or Glomax Discover (Promega). Results were reported as the percentage of viable cells with respect to untreated cells for two independent experiments.

### Luciferase reporter assay

Dual-luciferase plasmid pAsLuc(Fire)-HTLV-Luc(Reni) (Arpin-André *et al*, 2014) was kindly provided by Prof. Jean-Michel Mesnard from the Institut de Recherche en Infectiologie de Montpellier. The plasmid (500 ng) was transfected into HEK293T using Lipofectamine 3000 (ThermoFisher Scientific, #L3000015). 2 h post transfection, cells were treated with increasing concentrations of PDS. The LTR antisense promoter activity was assessed 24 h post-treatment through the firefly luciferase signal, measured by the Dual-Glo® Luciferase Assay System (Promega, #E2920) following the manufacturer’s instructions. The luciferase signal intensity was measured by Varioskan LUX Multimode Microplate Reader (ThermoFisher Scientific) and normalized to the total protein content, determined by the Pierce™ BCA Protein Assay Kit (ThermoFisher Scientific, # A55864). Statistical analysis was performed on Prism (version 10.0.3).

### Chromatin fixation and shearing

3×10^6^ MT-2 cells were fixed with 1% formaldehyde (ThermoFisher Scientific, #28906) for 10 min at 37°C shaking. Fixation was quenched by the addition of 125 mM glycine and incubated for 5 min. Cells were centrifuged for 5 min at 3500 rpm and washed twice with 10% FBS in PBS and centrifuged. Pellets were resuspended in 300 μL immunoprecipitation buffer (50 mM Hepes-KOH pH 7.5, 150 mM NaCl, 1 mM EDTA, 1% Triton X-100, 0.1% sodium deoxycholate, 0.1% SDS) and sonicated on Bioruptor Plus (Diagenode) at 4°C. 40 sonication cycles 30 sec on / 30 sec off were performed and fragments size was checked using 2100 Bioanalyzer system or TapeStation System (Agilent). Fixed chromatin samples were stored at −80 °C.

### BG4-ChIP-qPCR

G4 ChIP experiments were performed as previously described (Nicoletto *et al*, 2024), on three samples: INPUT, IP, Mock. 500 ng fixed chromatin for each sample was diluted in blocking buffer (25 mM HEPES pH 7.5, 10.5 mM NaCl, 110 mM KCl, 1 mM MgCl2, 1% BSA). Samples were incubated with 2 μL of RNase A (10 mg/mL ThermoFisher Scientific) at 37°C for 20 min at 800 rpm. After RNase A treatment, INPUT sample was stored on ice until the elution step. 250 ng of BG4 antibody (Maurizio *et al*, 2024) was added to IP sample. IP and MOCK were then incubated 1 h at 16°C, 1200 rpm. At the same time, anti-FLAG M2 magnetic beads (Merck, # M8823) were washed three times with blocking buffer, resuspended in the same buffer and incubated at 16°C, 1200 rpm. Then, 50 μL of magnetic beads were added to IP and MOCK samples, respectively, and incubated for 2 h at 16°C, 1200 rpm. Samples were then washed three times with ice-cold wash buffer (100 mM KCl, 0.1% Tween 20, and 10 mM Tris pH 7.4). Two additional washes were performed at 37°C, 1200 rpm for 10 min each. Elution was performed by adding TE buffer to the beads. Samples were treated with RNase A and proteinase K and DNA decrosslinking was obtained by incubating samples 8 h at 65°C. Samples were purified using MinElute PCR purification kit (QIAGEN, #28004) and qPCR was performed using LightCycler® 480 SYBR Green I Master (Roche, #04707516001) on LightCycler® 480 (Roche) quantitative PCR machine. Primers used for ChIP-qPCR are listed in Appendix Table S1. Statistical analysis was performed on Prism (version 10.0.3).

### SP1-ChIP-qPCR

The ChIP assay targeting SP1 was performed on 3.5 million MT-2 cells, either untreated or treated with 0.5 µM PDS for 24 h, according to previous procedures (Tosoni *et al*, 2025). Pre-cleared, sonicated chromatin was subjected to immunoprecipitation using an anti-SP1 antibody (Merck Millipore, #17-601) or a rabbit anti-mouse IgG antibody (Merck Millipore, #06-371) in immunoprecipitation buffer (100 mM Tris-HCl (pH 8.0), 100 mM NaCl, 5 mM EDTA, 0.3% sodium dodecyl sulfate, and 1.7% Triton X-100) overnight at 4°C. Then, the immunocomplexes were incubated with a protein A Dynabeads (ThermoFisher) for 4 h at 4°C and washed using four different buffers: wash buffer 1 (20 mM Tris-HCl (pH 8.0), 150 mM NaCl, 5 mM EDTA, 1% Triton X-100, 0.2% SDS, and 5% sucrose), wash buffer 2 (50 mM HEPES-NaOH (pH 7.5), 500 mM NaCl, 1 mM EDTA, 0.1% sodium deoxycholate, and Triton X-100), wash buffer 3 (10 mM Tris-HCl (pH 8.0), 250 mM LiCl, 1 mM EDTA, 0.5% sodium deoxycholate, and 0.5% IGEPAL), and wash buffer 4 (10 mM Tris-HCl (pH 8.0) and 1 mM EDTA). The chromatin fraction bound to Dynabeads was eluted in elution buffer (10 mM Tris-HCl (pH 8.0), 1 mM EDTA, and 1% SDS), and decrosslinked at 65°C for 4 h. Samples were subsequently treated with RNase A for 30 min at 37°C and proteinase K for 1 h at 55°C. DNA was purified using the MinElute PCR Purification Kit (Qiagen, Hilden, Germany), following the manufacturer’s instructions. qPCR was performed using SYBR™ Green PCR Master Mix (ThermoFisher) in QuantStudio™ 3 Real-Time PCR System (ThermoFisher) to measure the relative amounts of ChIP DNA relative to inputs. Primers used for ChIP-qPCR are listed in Appendix Table S1. Statistical analysis was performed on Prism (version 10.0.3).

### RT-PCR

MT-2 cells were seeded in 12-well plate for 24 h and treated with the indicated amounts of PDS. Total RNA was extracted 24 h post treatment using GeneJET RNA Purification Kit (ThermoFisher Scientific, #K0731) following manufacturer’s instructions and subjected to DNase digestion with TURBO DNA-free Kit (ThermoFisher Scientific, #AM1907) to remove genomic DNA contamination. Total RNA (250 ng) was reverse transcribed by TaqMan™ Reverse Transcription Reagents kit (ThermoFisher Scientific, #N8080234) using the oligo (dT16) to specifically amplify the total messenger RNA. The resultant complementary DNA was then amplified using SYBR™ Green PCR Master Mix (ThermoFisher Scientific, #4309155) in the presence of gene-specific primer pairs targeting HBZ and GAPDH regions (Appendix Table S1). Statistical analysis was performed on Prism (version 10.0.3).

## Acknowledgments

This work was supported by ESCMID Research Grant 2021 to E.R and by grants to S.N.R. from the European Research Council (ERC Consolidator 615879) and the Bill and Melinda Gates Foundation (OPP1035881 and OPP1097238).

We thank the BIONIC group at the University of Padua for helpful discussion.

## CRediT authorship contribution statement

Emanuela Ruggiero: Writing – review & editing, Writing – original draft, Methodology, Investigation, Formal analysis, Funding acquisition. Irene Zanin: Investigation, Formal analysis. Beatrice Tosoni: Investigation, Formal analysis. Sara N. Richter: Writing – review & editing, Supervision, Funding acquisition, Conceptualization.

## Supplementary Information

Appendix

